# Divergent methyl-coenzyme M reductase genes in a deep-subseafloor Archaeoglobi

**DOI:** 10.1101/390617

**Authors:** Joel A. Boyd, Sean P. Jungbluth, Andy O. Leu, Paul N. Evans, Ben J. Woodcroft, Grayson L. Chadwick, Victoria J. Orphan, Jan P. Amend, Michael S. Rappé, Gene W. Tyson

## Abstract

The methyl-coenzyme M reductase (MCR) complex is a key enzyme in archaeal methane generation and has recently been proposed to also be involved in the oxidation of short-chain hydrocarbons including methane, butane and potentially propane. The number of archaeal clades encoding the MCR complex continues to grow, suggesting that this complex was inherited from an ancient ancestor, or has undergone extensive horizontal gene transfer. Expanding the representation of MCR-encoding lineages through metagenomic approaches will help resolve the evolutionary history of this complex. Here, a near-complete Archaeoglobi metagenome-assembled genome (MAG; rG16) was recovered from the deep subseafloor along the Juan de Fuca Ridge flank that encodes two divergent McrABG operons similar to those found in *Candidatus* Bathyarchaeota and *Candidatus* Syntrophoarchaeum MAGs. rG16 is basal to members of the class Archaeoglobi, and encodes the genes for β-oxidation, potentially allowing an alkanotrophic metabolism similar to that proposed for *Ca.* Syntrophoarchaeum. rG16 also encodes a respiratory electron transport chain that can potentially utilize nitrate, iron, and sulfur compounds as electron acceptors. As the first Archaeoglobi with the MCR complex, rG16 extends our understanding of the evolution and distribution of novel MCR encoding lineages among the Archaea.

## Introduction

The methyl-coenzyme M reductase (MCR) complex is a key component of methane metabolism, and until recently had only been found within the Euryarchaeota (*Methanococcales, Methanopyrales, Methanobacteriales, Methanomicrobiales, Methanocellales, Methanosarcinales, Methanomassiliicoccales, Methanofastiosales, Methanoflorentales, Methanphagales* [ANME-1] and *Methanonatronarchaeia*). However, recent genome-centric metagenomic studies have led to the discovery of genomes encoding divergent MCR complexes within the *Candidatus* Bathyarchaeota and *Candidatus* Verstraetearchaeota^1,2^. Originally, the novel MCR-encoding *Ca.* Bathyarchaeota and Verstaetearchaeota were inferred to be capable of hydrogenotrophic and methylotrophic methanogenesis, respectively. Intriguingly, the *Ca.* Bathyarchaeota also appeared to be capable of producing energy through peptide fermentation and β-oxidation, unusual among MCR-encoding microorganisms. More recently a euryarchaeotal lineage, *Candidatus* Syntrophoarchaeum, was found to encode *Ca.* Bathyarchaeota-like MCR homologs and experimentally demonstrated to activate butane for oxidation via modified β-oxidation and Wood-Ljungdahl (WL) pathways^3^. The similarity in the MCR complexes and inferred metabolism of the *Ca.* Bathyarchaeota and *Ca.* Syntrophoarchaeum suggest that the *Ca.* Bathyarchaeota may also oxidise short hydrocarbons. Both organisms are confined to anoxic, hydrocarbon-rich habitats^1–3^, where abiotically produced short alkanes are abundant and likely to be utilized as carbon and energy sources. The increased number of archaeal lineages encoding the MCR complex and their metabolic flexibility suggests that these microorganisms may have a greater impact on carbon cycling than originally suspected.

The similarity of the MCR complexes encoded by *Ca.* Bathyarchaeota and *Ca.* Syntrophoarchaeum is incongruent with their large phylogenetic distance in the genome tree, suggesting that these genes were acquired via horizontal gene transfer (HGT)^4,5^. Both scenarios indicate that further diversity of divergent MCR-encoding lineages remains to be discovered^5^, which has been supported by gene-centric metagenomic analyses of deep-sea and terrestrial hydrothermal environments^6,7^. Expanding the genomic representation of novel MCR-encoding lineages by targeting these environments using genome-centric metagenomic approaches will help resolve the evolutionary history of the complex and expand the diversity of lineages known to be involved in hydrocarbon cycling.

Archaeoglobi is a class of thermophilic Archaea belonging to the Euryarchaeota that are abundant in subsurface hydrothermal environments, where they likely play a role in carbon and nutrient cycling^8,9^. The Archaeoglobi are split into three genera: *Archaeoglobus*, which are all heterotrophic or chemolithotrophic sulfate reducers^10–19^, and *Geoglobus* and *Ferroglobus*, which reduce both nitrate and ferric iron^20–22^. Pure cultures of Archaeoglobus have been shown to be capable of alkane oxidation^23,24^, and based on their shared metabolic features with methanogens^25–28^ and proximity to methanogens in the genome tree, are suggested to have an ancestor capable of methanogenesis. However, there are currently no representatives of the Archaeoglobi known to encode the MCR complex, likely a result of poor genomic representation caused by their extreme habitats that are difficult to sample.

Borehole observatories installed on the flank of the Juan de Fuca Ridge in the Pacific Ocean provide pristine fluids from the subseafloor igneous basement aquifer^29^. Previous metagenomic studies on samples collected from these borehole observatories revealed a distinct microbial community, including a number of novel Archaeoglobi^30,31^. Here, we characterise metagenome assembled genomes (MAGs) from igneous basement fluid samples from the boreholes^31^, focusing on a genome within a novel family that encodes two divergent copies of the *mcrABG* operon. Metabolic reconstruction revealed that the novel Archaeoglobi is potentially capable of hydrocarbon oxidation, amino acid fermentation, and can utilize multiple electron acceptors. Phylogenetic analyses support a horizontal gene transfer hypothesis for the distribution of novel MCR complex among the Archaea, and provides insight into the evolution of the Archaeoglobi.

## Results and discussion

To investigate the novel microbial diversity within Juan de Fuca Ridge flank boreholes, a metagenome (45.4 Gbp total raw reads) was from two wells were generated, assembled, and binned. Two of the 98 MAGs (rG3 and rG16) were found to encode full-length copies of the alpha subunit of the MCR complex (*mcrA*). rG3 encodes two McrA homologs with high sequence similarity to *Methanothermococcus thermolithotrophicus* (>99% amino acid identity, AAI). In contrast, rG16 (**Supplementary Note 1**; **Supplementary Figure 1**) encodes two divergent McrAs that were most similar to *Ca.* Syntrophoarchaeum caldarius (52% AAI) and *Ca.* Syntrophoarchaeum butanivorans (56% AAI). Based on 228 Euryarchaeota-specific marker genes, rG16 was estimated to be nearly complete (99.84%) with low contamination (1.96%), and a genome size of ~2.13 Mbp. Annotation of the 2,305 proteins encoded by the rG16 genome revealed all subunits of the MCR complex, including two copies of the *mcrC* subunit and an ancillary *mcrD* subunit, all of which have highest sequence similarity to homologs within *Ca.* Syntrophoarchaeum.

Phylogenetic analysis of the two MCR complexes in rG16 revealed that they branched with high bootstrap support to the divergent MCRs from *Ca.* Syntrophoarchaeum and *Ca.* Bathyarchaeota (**Figure 1A; Supplementary Figure 2 - 3**). Notably, the average branch length within the divergent McrA clade was double (1.05 ± 0.24 substitutions per site) that of traditional hydrogenotrophic, acetoclastic and H_2_-dependent methylotrophic methanogens (0.46 ± 0.10 substitutions per site), suggesting an accelerated rate of evolution following duplication or HGT^32^. To determine the taxonomy of rG16, a genome tree was constructed from a concatenated alignment of 122 archaeal single copy marker genes. rG16 was positioned basal to other members within the class Archaeoglobi with strong bootstrap support (**Figure 1B**; **Supplementary Figure 4**), including other Archaeoglobi previously recovered from the Juan de Fuca Ridge^31^. Phylogenetic analysis of the partial 16S rRNA gene (904 bp) confirmed the position of rG16 within the Archaeoglobi (**Supplementary Figure 5**). The average AAI between rG16 and other Archaeoglobi recovered from the Juan de Fuca Ridge (54.2% ± 0.7 AAI; **Supplementary Figure 6**) and relative evolutionary divergence^33,34^, suggest it is the first representative of a novel family within the Archaeoglobi^35^. The incongruencies between the genome tree and MCR phylogenies for rG16, *Ca.* Syntrophoarchaeum, and the *Ca.* Bathyarchaeota are most parsimoniously explained by HGT of the MCR.

**Figure 1.**
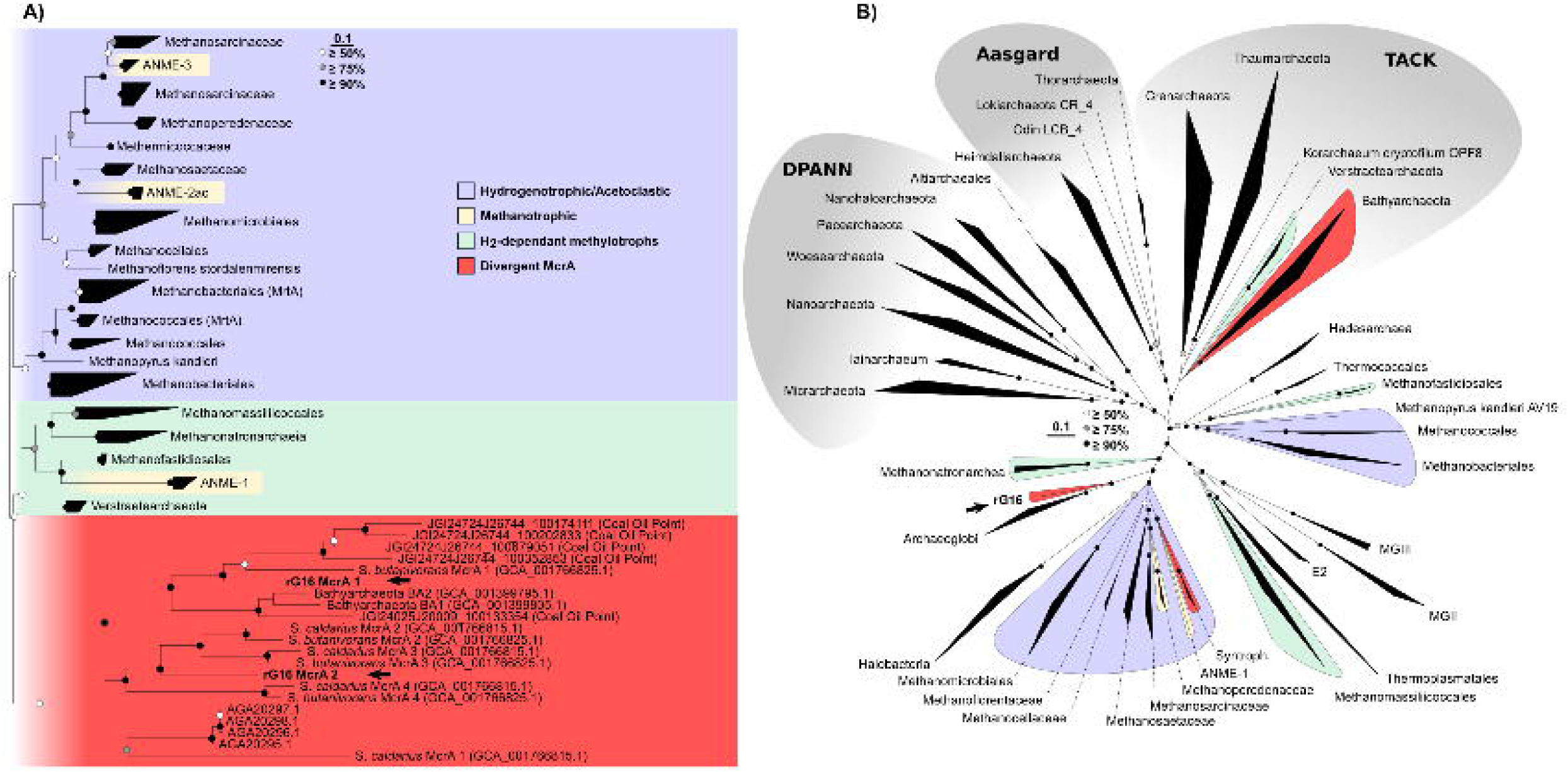
Maximum-likelihood trees of **A)** McrA proteins, and **B)** a concatenated alignment of 122 single copy archaeal marker genes from high quality archaeal RefSeq genomes (release 80). Hydrogenotrophic and acetoclastic methanogens are shaded blue, H_2_-dependent methylotrophic methanogens are shaded green, known/putative methane oxidisers are shaded yellow, and lineages encoding divergent McrAs are shaded red. Bootstrap support was generated from 100 replicates, and white, gray and black nodes represent ≥50%, ≥75% and ≥90% support, respectively.

**Figure 4.**
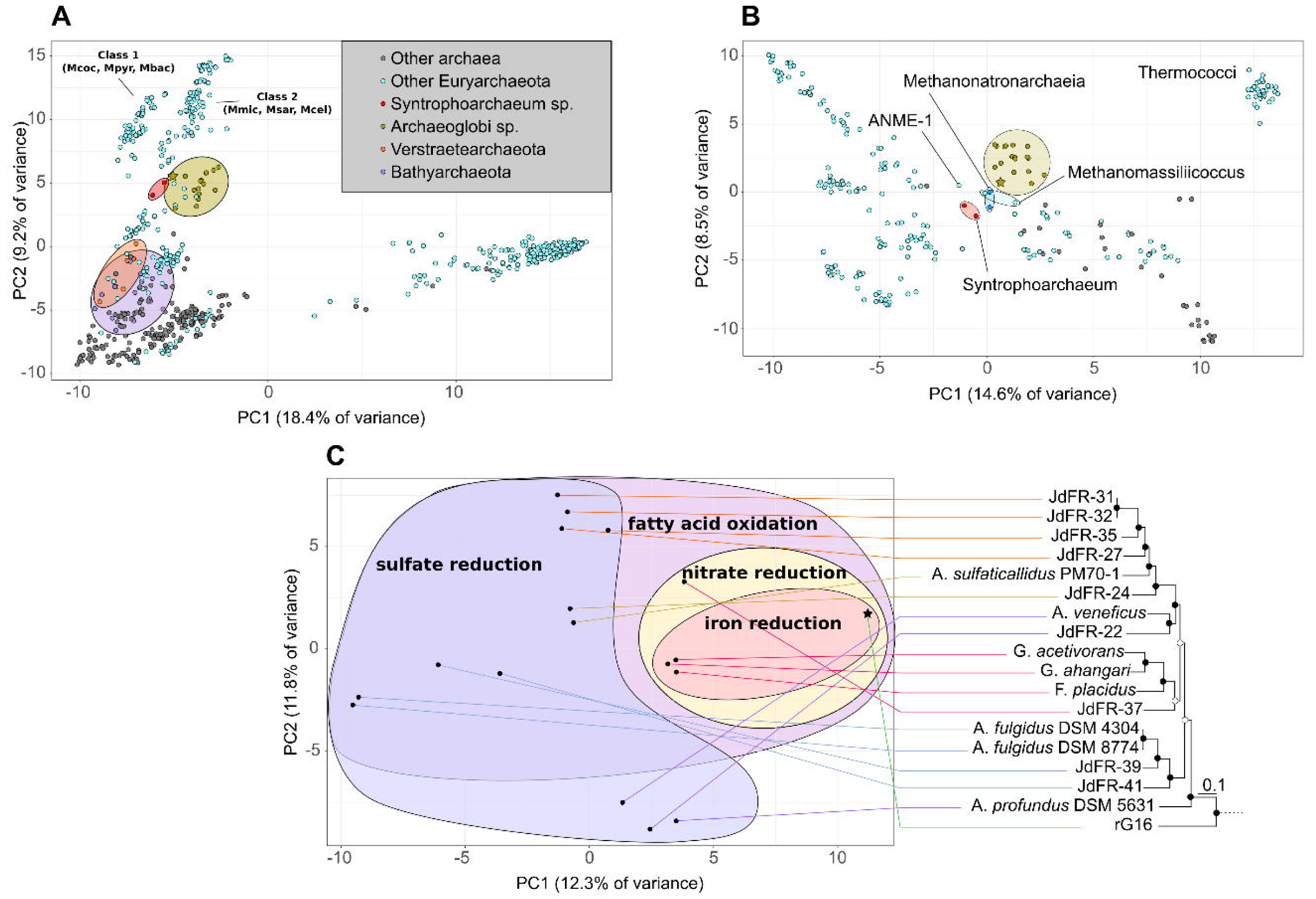
Comparative genomics of the rG16 genome. **A)** PCA of the presence/absence of KEGG Orthologous (KO) genes in all archaea, **B)** within the Euryarchaeota, with the exception of the Haloarchaea, and **C)** within the Archaeoglobi. The phylogeny of the Archaeoglobi within the maximum-likelihood tree of the 122 archaeal marker genes within **Figure 1B** is also shown.

Metabolic reconstruction of the rG16 MAG highlighted the potential for diverse metabolic capabilities, including amino acid fermentation and short chain alkane oxidation using a wide variety of electron acceptors (**Figure 2**). rG16 encodes a complete Wood-Ljungdahl pathway, and consistent with the *Ca.* Bathyarchaeota and *Ca.* Verstraetearchaeota, encodes 5 copies of the H subunit of the methyltetrahydromethanopterin (H_4_MPT): coenzyme M methyltransferase complex (*mtrH*), each co-located with predicted di- and tri-methylamine corrinoid proteins (**Supplementary Table 1**). This suggests that rG16 encodes a diverse range of methyltransferases, but is unlikely to conserve energy via methane oxidation or hydrogenotrophic methanogenesis^26^. The WL pathway may be used for oxidation of acetyl-CoA as previously observed in heterotrophic Archaeoglobales isolates^12^. *Ca.* Syntrophoarchaeum caldarius and *Ca.* Syntrophoarchaeum butanivorans have been inferred to oxidise alkanes activated by the MCR complex, putatively via the β-oxidation and WL pathways^3^. rG16’s two copies of McrA share catalytic residues with *Ca.* Syntrophoarchaeum homologs (**Supplementary Figure 7**), and encodes β-oxidation and methyltransferase enzymes that would allow short alkane oxidation (**Figure 2**). However, unlike *Ca.* Syntrophoarchaeum, the presence of a short chain acyl-CoA and butyryl-CoA dehydrogenase (*acd* and *bcd*, respectively), and a long-chain acyl-CoA synthetase (*fadD*) may allow rG16 to oxidise long chain fatty acids (**Figure 2**). The energy for hydrocarbon activation may be produced via either a soluble or membrane bound heterodisulfide reductase (*hdrABC*, *hdrDE*, respectively), both of which are encoded by the rG16 genome. While no *mvhAG* subunit was encoded by rG16, the C-terminal of *hdrA* is fused to the *mvhD* subunit as previously observed in *Methanosarcina acetivorans^36^* suggesting it plays a similar role in disulfide reduction. A further seven putative *hdrD* subunits co-located with flavin adenine dinucleotide-containing dehydrogenases (*glcD*) potentially oxidise coenzyme M (CoM-SH) and coenzyme B (CoB-SH) as proposed for the *Ca.* Verstraetearchaeota and *Ca.* Bathyarchaeota (**Figure 2**)^1,2^. While common to the Archaeoglobi, the membrane bound HdrDE has yet to be observed in *Ca.* Bathyarchaeota and *Ca.* Syntrophoarchaeum. The functional redundancy of *hdrABC*/*DE* has also been observed in *Archaeoglobus profundus^37^* where they were suggested to play a role in sulfur metabolism. However, without the dissimilatory sulfate reductase (*dsrAB*) gene, their role in rG16 remains unclear.

**Figure 2.**
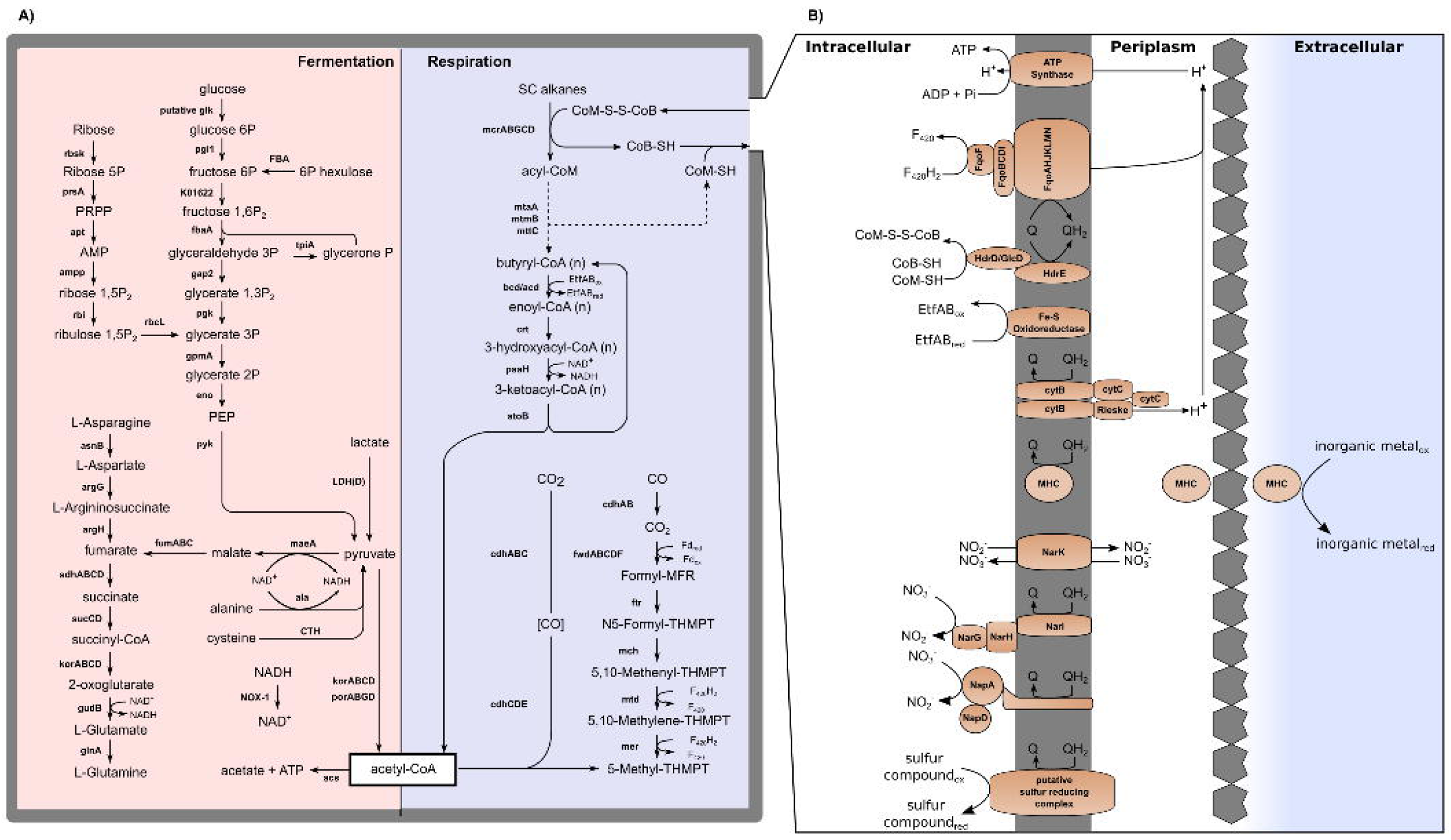
Metabolic reconstruction of the rG16 genome. **A)** Respiratory and fermentative pathways are shaded blue and red, respectively. **B)** Proposed CoM-S-S-CoB disulfide regeneration and membrane energetics of rG16 are shown. Genes associated with the pathways shown can be found in **Supplementary Table 1**.

Alkane oxidation can be energetically favourable when coupled to an electron acceptor such as sulfate^38^, nitrate^39^, nitrite^40^ and metal oxide^41^ reduction, or transferred to a syntrophic partner via direct interspecies electron transfer (DIET)^42^. Similar to the iron metabolising *Geoglobus^22,43,44^* and *Ferroglobus^45^* within the Archaeoglobi, rG16 does not encode *dsrAB*, but was found to encode 10 multi-haem c-type cytochromes (MHCs) with 4 - 31 haem binding motifs that may facilitate iron reduction^8,43,44,46^, or DIET as proposed in anaerobic methanotrophic archaea^42^. To compare the multi-haem cytochrome profile of rG16 with other Bacteria and Archaea, a network analysis was conducted on genomes from NCBI’s RefSeq database. Each MHC encoded by rG16 shared high sequence similarity to homologs encoded by archaeal (e.g. *Ferroglobus placidus*, *Geoglobus acetivorans*) and bacterial (e.g. *Ferrimonas* and *Shewanella*) iron reducers, *Ca.* Methanoperedens nitroreducens, and *Ca.* Syntrophoarchaeum (**Figure 3**). Four multi-haem cytochromes similar to *Methanoperedens*, *Ferroglobus* and *Geoglobus* homologs are organised into three contiguous operons (**scaffold 8; Figure 3**) encoding membrane-bound, redox-active complexes, including a bc1-like complex with a Rieske iron sulfur protein and cytochrome b that may generate a proton gradient with a Q-cycle (**scaffold 8, ORF 202 - 207; Figure 3**), and two complexes associated with the transfer of electrons to the membrane (**scaffold 8, ORF 212-215, ORF 208 - 211**). One complex includes an enzyme with two haem binding domains that is conserved among iron metabolising *Geoglobus* and *Ferroglobus* (**ORF 212**). Intriguingly, rG16 encodes an operon of three MHCs (**scaffold 16; ORF 2 - 4; Figure 3**), two of which are specific to Archaeoglobi, *Methanoperedenaceae*, and *Ca.* Syntrophoarchaeum, suggesting these MHCs play a specific role in alkane oxidisers (**Figure 3**). The final gene in this operon is homologous to a large C-type cytochrome also found in *Geoglobus acetivorans* SBH6^T^ that is a possible genomic determinant of iron reduction^21^. rG16 also encodes a *narGHJIK* (**Figure 3**), which may allow alkane oxidation coupled to nitrate reduction^39^. A further two operons encode sulfur reductase-like complexes that were previously only found in the hyperthermophile *Aquifex aeolicus^47^*, and shown to allow tetrathionate, polysulfide and elemental sulfur to be used as terminal electron acceptors (**Scaffold 6, ORF 76 - 80; Scaffold 4, ORF 33 - 36; Figure 3**). The potential to use partially oxidized forms of sulfur, nitrogen, and iron as electron acceptors suggests that rG16 can use different electron sinks depending on environmental conditions^48^.

**Figure 3.**
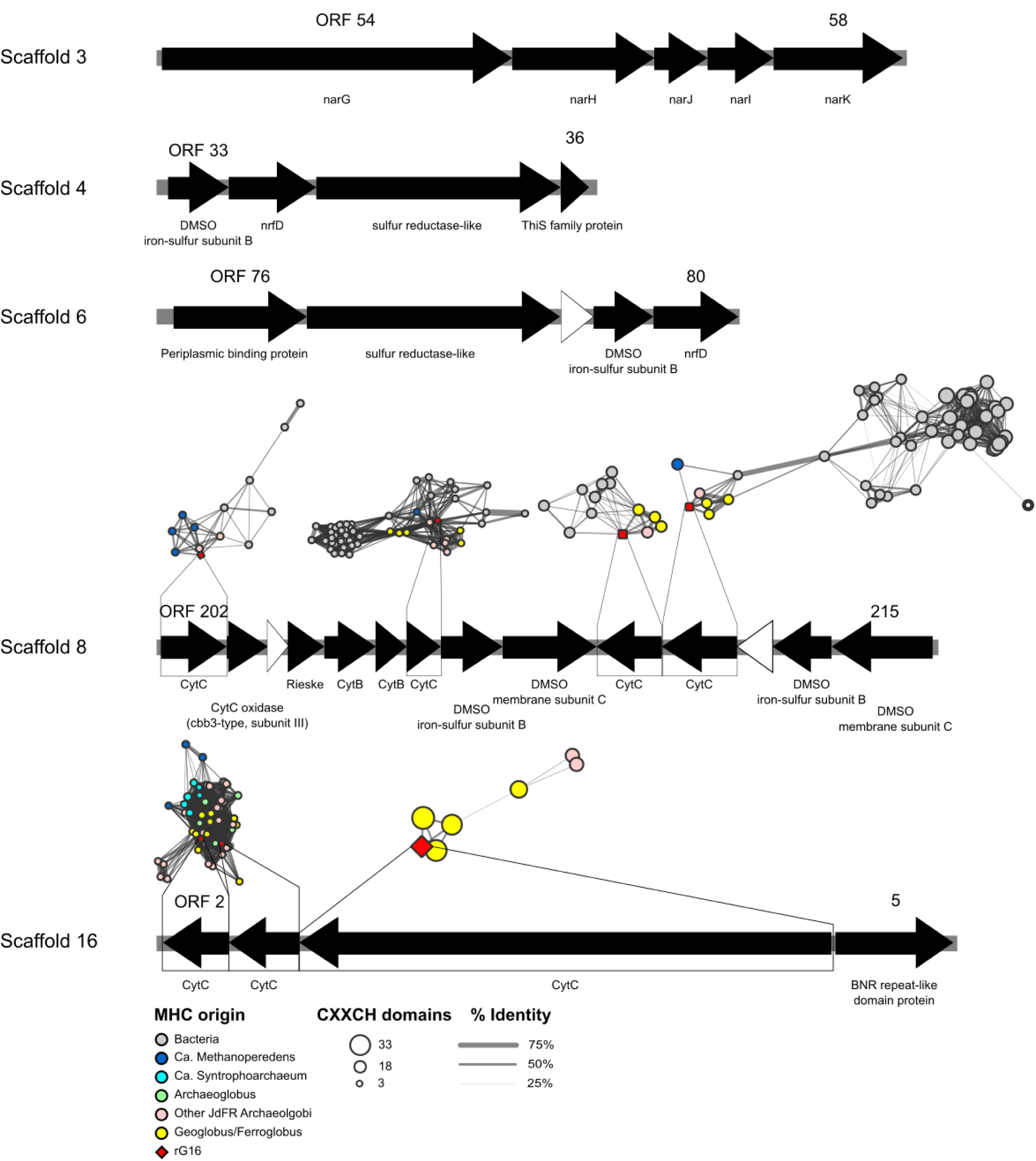
Operons encoding redox active complexes within rG16. White arrows represent hypothetical proteins. Networks are of MHCs and represent clusters of related proteins and their organism of origin.

rG16 putatively generates ATP via anaerobic respiration of short chain alkanes. A proton motive force is generated by the F_420_H_2_:quinone oxidoreductase, and ATP is generated by an archaeal type ATP synthase (**Figure 2, Box 1**). However, rG16 is also facultative fermenter, with the ability to ferment via a number of pathways (**Figure 2**). While no glucose transporters could be identified, glucose can be fermented via glycolysis, producing acetyl-CoA via pyruvate-ferredoxin oxidoreductase (*por*) or 2-oxoglutarate ferredoxin oxidoreductase (*kor*). rG16 also encodes genes to ferment organic acids such as lactate via lactate dehydrogenase (*ldh*) and succinate, fumarate and malate via a partial citric acid cycle (**Figure 2**). A type II/III ribulose 1,5-bisphosphate carboxylase/oxygenase homologous to *Ca.* Bathyarchaeota, *Ca.* Verstraetearchaeota, and Altiarchaeales may play a role in nucleotide catabolism^1,49^ (Archaeal type RuBisCO; **Supplementary Figure 10**). A number of amino acid transporters (**Branched chain amino acid transporters; scaffold 2, ORF 137 - 141, scaffold 4, ORF 1 - 5; scaffold 20, 82 - 84**) and peptidases (tetrahedral aminopeptidase, peptidase family M50, Xaa-Pro dipeptidase, methionine aminopeptidase) indicate that rG16 can also ferment peptides (**Figure 2**). Pathways for the fermentation of glutamate, glutamine, alanine, cysteine, aspartate and asparagine exist, as well as a number of aminotransferases (*aspB*, *hisC*, *cobD*) and two copies of *por*, which is involved in the fermentation of aromatic amino acids in other hyperthermophiles^50^. rG16 also encodes a benzoyl-CoA reductase complex (*bcrABCD*) indicating the potential to degrade aromatic compounds. The acetyl-CoA generated via glycolysis and via fermentation of organic and amino acids can be used for substrate level-phosphorylation using acetyl-CoA synthetase (*acs*) or acetate-CoA ligase (*acd*). The various fermentative strategies used by rG16 suggest that it is adapted to a fluctuating availability of organic compounds.

To compare the metabolic capabilities of rG16 with publically available archaeal genomes from RefSeq and GenBank, a global analyses of KEGG Orthologous (KO) genes was conducted. The KO profile of rG16 was most similar to other members of the Archaeoglobi, *Methanomassiliicoccales* ANME-1, *Candidatus* Syntrophoarchaeales and *Methanonatronarchaeia sp.*, but distant from the *Ca.* Bathyarchaeota and *Ca.* Verstraetearchaeota (**Figure 4A, B**). To further examine the shared genomic content of novel MCR encoding lineages, orthologous clusters (OCs) were generated using proteinortho^51^. Within the Archaeoglobi, 134 OCs were unique to rG16, mapping to 71 KOs primarily associated with carbon metabolism (**Supplementary Table 1**). The few clusters that were unique to the rG16, *Ca.* Bathyarchaeota and *Ca.* Syntrophoarchaeum (seven OCs) were limited to subunits from the MCR complex, a putative methanogenesis marker (TIGR03275), and a gene associated with cobalamin biosynthesis (cob(I)alamin adenosyltransferase; *cobA*; **Supplementary Table 1**). A further 25 OCs were specific only to rG16 and *Ca.* Syntrophoarchaeum, including four further methanogenesis markers (TIGR03271, TIGR03291, TIGR03268, TIGR03282), a sugar-specific transcriptional regulator, and a class II fumarate hydratase. Many OCs shared between rG16 and *Ca.* Syntrophoarchaeum were annotated as ‘hypothetical protein’, suggesting much of the metabolic similarities of novel hydrocarbon metabolisers have yet to be functionally characterised.

To explore the evolutionary history of the MCR complex, the gene phylogeny of the McrA subunit was compared with the archaeal genome phylogeny (**Supplementary Figure 9**). Traditional euryarchaeal methanogens are largely congruent with the branching order of the genome tree (**Supplementary Figure 9**), suggesting that the evolutionary history of the McrA largely follows vertical inheritance. However, the monophyletic H_2_-dependent methylotroph and *Ca.* Bathyarchaeota/*Ca.* Syntrophoarchaeum/rG16 clades are highly paraphyletic in the genome tree (**Supplementary Figure 9**). *Ca.* Bathyarchaeota and Archaeoglobi McrA cluster with different homologs of the *Ca.* Syntrophoarchaeum (**Figure 1A**), a phylogenetic pattern most parsimoniously explained by HGT (**Supplementary Figure 9)**. The basal phylogenetic position of rG16 and divergence to other *Ca.* Syntrophoarcheum MCR would suggest this event did not occur in recent evolutionary history, supported by the lack of a divergent GC or kmer profile surrounding the gene context of the rG16’s MCRs^52^ (**Supplementary Figure 10 - 11**). Given the metabolic similarities between the Archaeoglobi and methanogens^26^, it has been hypothesised that the last common ancestor (LCA) of the Archaeoglobi encoded the MCR complex, which was subsequently lost following HGT of the *dsrAB* gene from the Bacteria^53,54^. While it is unclear whether the LCA of the Archaeoglobi encoded a conventional or divergent MCR complex, three scenarios explain the current distribution of *dsrAB* and the MCR complex in this lineage: i) HGT of *dsrAB* into the LCA of the Archaeoglobi, followed by loss of this metabolism after acquisition of the divergent MCR complex in rG16 (**Supplementary Figure 12A**), ii) HGT of the divergent MCR complex into the LCA of the Archaeoglobi, followed by loss of the complex after the acquisition of *dsrAB* in the Archaeoglobaceae (**Supplementary Figure 12B**), or iii) the separate acquisition of the divergent MCR complex and *dsrAB* by rG16 and the Archaeoglobaceae (**Supplementary Figure 12C**). Greater genomic representation and comparative genomics of the Archaeoglobi will clarify the evolutionary story, particularly if MCR-encoding lineages are found that fall within the Archaeoglobaceae. The evolutionary history of the MCR is highly complex, with evidence for both vertical inheritance and HGT (**Supplementary Figure 12**). It is likely that archaeal lineages encoding the divergent MCR complex will continue to be discovered, and with the genomic representation they add to public databases, their evolutionary history and metabolic role in the hydrothermal subsurface biosphere will become increasingly clear.

## Methods

### Metagenome generation, assembly and binning

Two metagenomes from crustal fluids of the JdFR flank were generated as described previously^31^. One sample (SRR3723048) that yielded the novel Archaeoglobi genome rG16 was selected for reassembly using metaSPAdes v3.9.0^55^ with default settings, using raw reads as input. Raw reads were mapped to the resulting assembly using BWA-MEM^56^ v0.7.12. Binning was conducted using MetaBAT v0.32.4^57^ using the --specific setting.

### Identification of MCR encoding genomes

Genomes generated by MetaBAT were searched with GraftM v0.11.1^6^ using an McrA-specific GraftM package (gpkg). The McrA gene tree was curated with NCBI taxonomy, with the Bathyarchaeota and Syntrophoarcaheum clade labeled as “divergent”. An evalue of 1e-50 was used to filter for full-length McrA genes.

### MAG quality control

rG16 was analysed using RefineM^58^ v0.0.23 to identify contigs with divergent tetranucleotide frequencies and GC content. A single 2,748 bp contig was removed due to divergent a GC, tetranucleotide and taxon profile (**Supplementary Note 1**). The remaining contigs were scaffolded with FinishM v0.0.7 roundup using default parameters (github.com/wwood/finishm). The completeness and contamination of the resulting bin was assessed using CheckM v1.0.8^59^ with default settings.

### Annotation of rG16

The rG16 MAG was annotated using EnrichM annotate (github.com/geronimp/enrichM). Briefly, EnrichM calls proteins from contigs using Prodigal v2.6.3^60^, and blasts them against UniRef100 using DIAMOND^61^ v0.9.22 to obtain KO annotations. Pfam-A^62^ (release 32) and TIGRFAM^63^ (release 15.0) Hidden Markov Models (HMMs) were run on the proteins using hmmer 3.1b^64^ to obtain Pfam and TIGRFAM annotations, respectively. Further manual curation was completed using NCBI BLAST and CD-Search^65^.

### Genome tree

Using GenomeTreeTK (https://github.com/dparks1134/GenomeTreeTk) v0.0.41, a genome tree of Archaea from NCBI’s RefSeq database (release 80) was created using a concatenated alignment of 122 archaea-specific single marker copy genes. Genomes <50% complete, and with >10% contamination as determined using CheckM were removed from the analysis. After alignment to HMMs constructed for each of the 122 marker genes, alignments were concatenated and genomes with <50% of the alignment were excluded from the analysis. Maximum likelihood trees were constructed using FastTree v2.1.9, and non-parametric bootstrapping was completed using GenomeTreeTK’s bootstrap function.

### 16S rRNA gene tree

Sequences classified as Archaeoglobi with a pintail score of 100 and an alignment and sequence quality of ≥ 80 were extracted from the SILVA database^66^ (version 132) and used as reference sequences for a 16S rRNA phylogenetic tree. The partial 16S rRNA sequence from rG16 was added to the database, sequences were aligned using ssu-align^67^ v0.1, and subsequently converted to fasta format using the convert from seqmagick v0.6.1 (fhcrc.github.io/seqmagick). Gapped regions in the alignment were removed with trimAl v1.2 using the --gappyout flag^68^. The Maximum likelihood tree was constructed using FastTreeMP^69^ with a generalized time-reversible model and --nt flags, bootstrapped with GenomeTreeTK (github.com/dparks1134/GenomeTreeTk), and visualised in ARB^70^ v6.0.6.

### Gene phylogenies (McrA, McrB, McrG, RuBisCo)

McrA, McrB, McrG and RuBisCo sequences were derived from the genomes used in the genome tree. Proteins from all genomes were called using Prodigal v2.6.3, and searched using hmmer v3.1b with Pfam Hidden Markov models (PF02240.15, MCR_gamma; PF02241.17, MCR_beta; PF02249.16, MCR_alpha; PF02745.14, MCR_alpha_N; PF00016.19, RuBisCO_large; PF02788.15, RuBisCO_large_N) with the --cut_tc flag to minimize false positives. For McrA and RuBisCo, both models needed to hit a sequence for it to be included in the analysis. For each gene, sequences were aligned using MAFFT-GINS-i v7.221 and filtered using trimAl with the --gappyout flag. A maximum likelihood tree was constructed using FastTreeMP with default parameters.

### Average amino acid identity

Average Amino acid identity was generated with CompareM v0.0.22 (https://github.com/dparks1134/CompareM) using the aai_wf with default parameters.

### Network analysis of MHCs

Proteins from the Archaea in NCBI’s RefSeq database (release 80) were searched with the Cytochrome C Pfam HMM (PF00034). Hits were filtered to have at least one of the Cytochrome C CXXCH domains using a custom script (fastacxxch.count.py, github.com/geronimp/HandyScripts/blob/master/99_random/fastacxxch.count.py). Closest matches for all resulting proteins were identified using DIAMOND with an evalue cutoff of 1e-20. The result was visualised in Cytoscape v3.2.0, removing clusters without a rG16 homolog.

### KO analysis

Proteins from the genomes were searched using DIAMOND blastp against UniRef100 with an evalue cutoff of 1e-05. For each protein, the KO annotations were derived from the top hit. The presence/absence of each KO in each genome was used as input to a Principal Component Analysis (PCA) using the prcomp function in R.

### Sliding window GC and tetranucleotide frequencies

A custom script (https://github.com/geronimp/window_sequence) was written to fragment the rG16 contigs into short sequences, in a sliding window. For each fragment, the percent GC and 4mer frequency was calculated using seqstat in the biosquid package v1.9g (packages.debian.org/sid/biosquid) and Jellyfish^71^ v2.2.6.

### Data visualisation

Figures were generated in R^72^ v3.0.1 using ggplot^73^ v1.0.0 and refined using Inkscape v0.91 (inkscape.org/en).

## Author Contributions

J.A.B. and S.P.J. contributed equally to this work. M.S.R., J.P.A., and G.W.T. designed the overall study and procured funding. J.A.B., S.P.J., M.S.R and G.W.T. designed and carried out experiments and analyses around specific microbial hypotheses. J.A.B, S.P.J and G.W.T. wrote the manuscript. All authors edited, reviewed and approved the final manuscript.

## Acknowledgements

We thank the captain and crew, A. Fisher, K. Becker, C. G. Wheat, and other members of the science teams on board R/V Atlantis cruise AT18-07. We also thank the pilots and crew of remote-operated vehicle Jason II. This research was supported by two grants from the National Science Foundation: Microbial Observatories (MCB06-04014 to M. S. R.), and the Science and Technology Center for Dark Energy Biosphere Investigations (C-DEBI; OCE-0939564 to J. P. A). This study used samples and data provided by the Integrated Ocean Drilling Program. This material is based upon work supported by the U.S. Department of Energy, Office of Science, Office of Biological and Environmental Research program under Award Number. Genomic Science Program of the United States Department of Energy Office of Biological and Environmental Research [DE-SC0016469, DE-SC0010580, DE-SC0016440]; University of Queensland Vice Chancellor Research Focused Fellowship (to G. W. T.); Australian Research Council (ARC) Postgraduate Award (to J. A. B.); ARC Discovery Early Career Researcher Award (DE-160100248 to B. J. W.); ARC DECRA (170100428 to P. N. E.).

## Data Availability

The datasets analysed during the current study are available in the NCBI SRA, accession number SRR3723048.

## Code Availability

All unpublished software used in this publication are available on github; sequence_windower (https://github.com/geronimp/window_sequence); EnrichM (github.com/geronimp/enrichM); GenomeTreeTK (github.com/dparks1134/GenomeTreeTk); CompareM v0.0.22 (https://github.com/dparks1134/CompareM); fastacxxch.count.py, github.com/geronimp/HandyScripts/blob/master/99_random/fastacxxch.count.py; FinishM v0.0.7 github.com/wwood/finishm.

## References

1. Evans, P. N. et al. Methane metabolism in the archaeal phylum Bathyarchaeota revealed by genome-centric metagenomics. Science 350, 434–438 (2015).

2. Vanwonterghem, I. et al. Methylotrophic methanogenesis discovered in the archaeal phylum Verstraetearchaeota. Nature Microbiology 1, 16170 (2016).

3. Laso-Pérez, R. et al. Thermophilic archaea activate butane via alkyl-coenzyme M formation. Nature539 396–401 (2016).

4. Adam, P. S., Borrel, G., Brochier-Armanet, C. & Gribaldo, S. The growing tree of Archaea: new perspectives on their diversity, evolution and ecology. ISME Journal 11, 2407–2425 (2017).

5. Evans, P., Boyd, J., Leu, A., Donovan, W. B. P. & Tyson, G. Phylogenetic diversity and metabolic capacity of mcr and mcr-like containing archaeal lineages. Submitted

6. Boyd, J. A., Woodcroft, B. J. & Tyson, G. W. GraftM: a tool for scalable, phylogenetically informed classification of genes within metagenomes. Nucleic Acids Research 46, e59–e59 (2018).

7. McKay, L. J., Hatzenpichler, R., Inskeep, W. P. & Fields, M. W. Occurrence and expression of novel methyl-coenzyme M reductase gene (mcrA) variants in hot spring sediments. Scientific Reports 7, 7252 (2017).

8. Brileya, K. & Reysenbach, A.-L. The Class Archaeoglobi. in The Prokaryotes (eds. Rosenberg, E., DeLong, E. F., Lory, S., Stackebrandt, E. & Thompson, F.) 15–23 (Springer Berlin Heidelberg, 2014).

9. Orcutt, B. N., Sylvan, J. B., Knab, N. J. & Edwards, K. J. Microbial ecology of the dark ocean above, at, and below the seafloor. Microbiol. Mol. Biol. Rev. 75, 361–422 (2011).

10. Birkeland, N.-K., Schönheit, P., Poghosyan, L., Fiebig, A. & Klenk, H.-P. Complete genome sequence analysis of Archaeoglobus fulgidus strain 7324 (DSM 8774), a hyperthermophilic archaeal sulfate reducer from a North Sea oil field. Standards in Genomic Sciences 12, 79 (2017).

11. Stetter, K. O. Archaeoglobus fulgidus gen. nov., sp. nov.: a new taxon of extremely thermophilic archaebacteria. Applied and Environmental Microbiology 10, 172–173 (1988).

12. Klenk, H. P. et al. The complete genome sequence of the hyperthermophilic, sulphate-reducing archaeon Archaeoglobus fulgidus. Nature 390, 364–370 (1997).

13. Burggraf, S., Jannasch, H. W., Nicolaus, B. & Stetter, K. O. Archaeoglobus profundus sp. nov., Represents a New Species within the Sulfate-reducing Archaebacteria. Systematic and Applied Microbiology 13, 24–28 (1990).

14. von Jan, M. et al. Complete genome sequence of Archaeoglobus profundus type strain (AV18). Standards in Genomic Sciences 2, 327–346 (2010).

15. Huber, H., Jannasch, H., Rachel, R., Fuchs, T. & Stetter, K. O. Archaeoglobus veneficus sp. nov., a Novel Facultative Chemolithoautotrophic Hyperthermophilic Sulfite Reducer, Isolated from Abyssal Black Smokers. Systematic and Applied Microbiology 20, 374–380 (1997).

16. Mori, K., Maruyama, A., Urabe, T., Suzuki, K.-I. & Hanada, S. Archaeoglobus infectus sp. nov., a novel thermophilic, chemolithoheterotrophic archaeon isolated from a deep-sea rock collected at Suiyo Seamount, Izu-Bonin Arc, western Pacific Ocean. International Journal of Systematic and Evolutionary Microbiology 58, 810–816 (2008).

17. Steinsbu, B. O. et al. Archaeoglobus sulfaticallidus sp. nov., a thermophilic and facultatively lithoautotrophic sulfate-reducer isolated from black rust exposed to hot ridge flank crustal fluids. International Journal of Systematic and Evolutionary Microbiology 60, 2745–2752 (2010).

18. Stokke, R., Hocking, W. P., Steinsbu, B. O. & Steen, I. H. Complete Genome Sequence of the Thermophilic and Facultatively Chemolithoautotrophic Sulfate Reducer Archaeoglobus sulfaticallidus Strain PM70-1T. Genome Announcements 1, (2013).

19. Stetter, K. O. et al. Hyperthermophilic archaea are thriving in deep North Sea and Alaskan oil reservoirs. Nature 365, 743 (1993).

20. Anderson, I. et al. Complete genome sequence of Ferroglobus placidus AEDII12DO. Standards in Genomic Sciences 5, 50–60 (2011).

21. Mardanov, A. et al. The Geoglobus acetivorans genome: Fe(III) reduction, acetate utilization, autotrophic growth, and degradation of aromatic compounds in a hyperthermophilic archaeon. Applied and Environmental Microbiology 81, 1003–1012 (2015).

22. Manzella, M. P. et al. The complete genome sequence and emendation of the hyperthermophilic, obligate iron-reducing archaeon ‘Geoglobus ahangari’ strain 234 T. Standards in Genomic Sciences 10, (2015).

23. Khelifi, N. et al. Anaerobic oxidation of long-chain n-alkanes by the hyperthermophilic sulfate-reducing archaeon, Archaeoglobus fulgidus. ISME Journal 8, 2153–2166 (2014).

24. Khelifi, N. et al. Anaerobic oxidation of fatty acids and alkenes by the hyperthermophilic sulfate-reducing archaeon Archaeoglobus fulgidus. Applied and Environmental Microbiology 76, 3057–3060 (2010).

25. Ney, B. et al. The methanogenic redox cofactor F420 is widely synthesized by aerobic soil bacteria. ISME Journal 11, 125–137 (2017).

26. Borrel, G., Adam, P. S. & Gribaldo, S. Methanogenesis and the Wood–Ljungdahl Pathway: An Ancient, Versatile, and Fragile Association. Genome Biology and Evolution 8, 1706–1711 (2016).

27. Vornolt, J., Kunow, J., Stetter, K. O. & Thauer, R. K. Enzymes and coenzymes of the carbon monoxide dehydrogenase pathway for autotrophic CO2 fixation in Archaeoglobus lithotrophicus and the lack of carbon monoxide dehydrogenase in the heterotrophic A. profundus. Archives of Microbiology 163, 112–118 (1995).

28. Bapteste, E., Brochier, C. & Boucher, Y. Higher-level classification of the Archaea: evolution of methanogenesis and methanogens. Archaea 1, 353–363 (2005).

29. Lin, H.-T., Cowen, J. P., Olson, E. J., Amend, J. P. & Lilley, M. D. Inorganic chemistry, gas compositions and dissolved organic carbon in fluids from sedimented young basaltic crust on the Juan de Fuca Ridge flanks. Geochimica et Cosmochimica Acta 85, 213–227 (2012).

30. Jungbluth, S. P., Bowers, R. M., Lin, H.-T., Cowen, J. P. & Rappé, M. S. Novel microbial assemblages inhabiting crustal fluids within mid-ocean ridge flank subsurface basalt. ISME Journal 10, 2033–2047 (2016).

31. Jungbluth, S. P., Amend, J. P. & Rappé, M. S. Metagenome sequencing and 98 microbial genomes from Juan de Fuca Ridge flank subsurface fluids. Scientific Data 4, 170037 (2017).

32. Petitjean, C., Makarova, K., Wolf, Y. & Koonin, E. Extreme Deviations from Expected Evolutionary Rates in Archaeal Protein Families. Genome Biology and Evolution 9, 2791–2811 (2017).

33. Chaumeil, P.-A., Hugenholtz, P. & Parks, D. H. GTDB-Tk: A toolkit to classify genomes with the Genome Taxonomy Database. in preparation (2018).

34. Parks, D. H., Chuvochina, M., Waite, D. W. & Rinke, C. A proposal for a standardized bacterial taxonomy based on genome phylogeny. bioRxiv (2018).

35. Konstantinidis, K. T. & Tiedje, J. M. Towards a genome-based taxonomy for prokaryotes. Journal of Bacteriology 187, 6258–6264 (2005).

36. Yan, Z., Wang, M. & Ferry, J. G. A Ferredoxin- and F420H2-Dependent, Electron-Bifurcating, Heterodisulfide Reductase with Homologs in the Domains Bacteria and Archaea. MBio 8, (2017).

37. Mander, G. J., Pierik, A. J., Huber, H. & Hedderich, R. Two distinct heterodisulfide reductase-like enzymes in the sulfate-reducing archaeon Archaeoglobus profundus. FEBS J. 271, 1106–1116 (2004).

38. Milucka, J. et al. Zero-valent sulphur is a key intermediate in marine methane oxidation. Nature 491,541–546 (2012).

39. Haroon, M. F. et al. Anaerobic oxidation of methane coupled to nitrate reduction in a novel archaeal lineage. Nature 500, 567–570 (2013).

40. Ettwig, K. F. et al. Nitrite-driven anaerobic methane oxidation by oxygenic bacteria. Nature 464, 543–548 (2010).

41. Ettwig, K. F. et al. Archaea catalyze iron-dependent anaerobic oxidation of methane. Proceedings of the National Academy of Sciences of the United States of America 113, 12792–12796 (2016).

42. McGlynn, S. E., Chadwick, G. L., Kempes, C. P. & Orphan, V. J. Single cell activity reveals direct electron transfer in methanotrophic consortia. Nature 526, 531–535 (2015).

43. Slobodkina, G., Kolganova, T., Querellou, J., Bonch-Osmolovskaya, E. & Slobodkin, A. Geoglobus acetivorans sp. nov., an iron (III)-reducing archaeon from a deep-sea hydrothermal vent. International Journal of Systematic and Evolutionary Microbiology 59, 2880–2883 (2009).

44. Kashefi, k. et al. Geoglobus ahangari gen. nov., sp. nov., a novel hyperthermophilic archaeon capable of oxidizing organic acids and growing autotrophically on hydrogen with Fe(III) serving as the sole electron acceptor. International Journal of Systematic and Evolutionary Microbiology 52, 719–728 (2002).

45. Hafenbradl, D. et al. Ferroglobus placidus gen. nov., sp. nov., a novel hyperthermophilic archaeum that oxidizes Fe2+ at neutral pH under anoxic conditions. Archives of Microbiology 166, 308–314 (1996).

46. Tor, J. M. & Lovley, D. R. Anaerobic degradation of aromatic compounds coupled to Fe(III) reduction by Ferroglobus placidus. Environmental Microbiology 3, 281–287 (2001).

47. Guiral, M. et al. A Membrane-bound Multienzyme, Hydrogen-oxidizing, and Sulfur-reducing Complex from the Hyperthermophilic Bacterium Aquifex aeolicus. The Journal of Biological Chemistry 280, 42004–42015 (2005).

48. Wankel, S. D. et al. Anaerobic methane oxidation in metalliferous hydrothermal sediments: influence on carbon flux and decoupling from sulfate reduction. Environmental Microbiology 14, 2726–2740 (2012).

49. Wrighton, K. C. et al. Metabolic interdependencies between phylogenetically novel fermenters and respiratory organisms in an unconfined aquifer. ISME Journal 8, 1452–1463 (2014).

50. Mai, X. & Adams, M. W. Indolepyruvate ferredoxin oxidoreductase from the hyperthermophilic archaeon Pyrococcus furiosus. A new enzyme involved in peptide fermentation. The Journal of Biological Chemistry 269, 16726–16732 (1994).

51. Lechner, M. et al. Proteinortho: Detection of (Co-)orthologs in large-scale analysis. BMCBioinformatics 12, 124 (2011).

52. Ravenhall, M., Škunca, N., Lassalle, F. & Dessimoz, C. Inferring horizontal gene transfer. PLoS Computational Biology 11, e1004095 (2015).

53. Klein, M. et al. Multiple lateral transfers of dissimilatory sulfite reductase genes between major lineages of sulfate-reducing prokaryotes. Journal of Bacteriology 183, 6028–6035 (2001).

54. Müller, A. L., Kjeldsen, K. U., Rattei, T., Pester, M. & Loy, A. Phylogenetic and environmental diversity of DsrAB-type dissimilatory (bi)sulfite reductases. ISME J. 9, 1152 (2014).

55. Nurk, S., Meleshko, D., Korobeynikov, A. & Pevzner, P. A. metaSPAdes: a new versatile metagenomic assembler. Genome Research 27, 824–834 (2017).

56. Li, H. & Durbin, R. Fast and accurate short read alignment with Burrows-Wheeler transform. Bioinformatics 25, 1754–1760 (2009).

57. Kang, D. D., Froula, J., Egan, R. & Wang, Z. MetaBAT, an efficient tool for accurately reconstructing single genomes from complex microbial communities. PeerJ 3, e1165 (2015).

58. Parks, D. H. et al. Recovery of nearly 8,000 metagenome-assembled genomes substantially expands the tree of life. Nature Microbiology 2, 1533–1542 (2017).

59. Parks, D. H., Imelfort, M., Skennerton, C. T., Hugenholtz, P. & Tyson, G. W. CheckM: assessing the quality of microbial genomes recovered from isolates, single cells, and metagenomes. Genome Research 25, 1043–1055 (2015).

60. Hyatt, D. et al. Prodigal: prokaryotic gene recognition and translation initiation site identification. BMC Bioinformatics 11, 119 (2010).

61. Buchfink, B., Xie, C. & Huson, D. H. Fast and sensitive protein alignment using DIAMOND. Nature Methods 12, 59–60 (2014).

62. Finn, R. D. et al. The Pfam protein families database: towards a more sustainable future. Nucleic Acids Research 44, D279–D285 (2015).

63. Selengut, J. D. et al. TIGRFAMs and Genome Properties: tools for the assignment of molecular function and biological process in prokaryotic genomes. Nucleic Acids Research 35, D260–4 (2007).

64. Eddy, S. R. Accelerated Profile HMM Searches. PLoS computational biology 7, e1002195 (2011).

65. Marchler-Bauer, A. & Bryant, S. H. CD-Search: protein domain annotations on the fly. Nucleic Acids Res. 32, W327–31 (2004).

66. Quast, C. et al. The SILVA ribosomal RNA gene database project: improved data processing and web-based tools. Nucleic Acids Research 41, D590–D596 (2013).

67. Nawrocki, E. P. & Eddy, S. R. ssu-align: a tool for structural alignment of SSU rRNA sequences. (2010).

68. Capella-Gutiérrez, S., Silla-Martínez, J. M. & Gabaldón, T. trimAl: a tool for automated alignment trimming in large-scale phylogenetic analyses. Bioinformatics 25, 1972–1973 (2009).

69. Price, M., Dehal, P. & Arkin, A. FastTree 2–approximately maximum-likelihood trees for large alignments. PLoS One 5, e9490 (2010).

70. Ludwig, W. et al. ARB: a software environment for sequence data. Nucleic Acids Research 32, 1363–1371 (2004).

71. Marçais, G. & Kingsford, C. A fast, lock-free approach for efficient parallel counting of occurrences of k-mers. Bioinformatics 27, 764–770 (2011).

72. Team, R. C. & Others. R: A language and environment for statistical computing. (2013).

73. Wickham, H. ggplot2: Elegant Graphics for Data Analysis. (Springer-Verlag New York, 2009).

